# A general inverse problem approach to neural function modeling in *Drosophila melanogaster*

**DOI:** 10.1101/2020.05.08.084962

**Authors:** Marco Stucchi

## Abstract

The development of models of brain function remains a complex problem given the difficulty of extracting organizational principles from observations on a variety of morphologically and physiologically different neurons. State of the art results in this modeling research have been obtained by a different route, by leveraging the power of deep learning. However, this approach takes advantage of neuroscientific knowledge only to a limited extent. Here, I adopt a perspective that aims at combining experimental data and optimization algorithms by framing this modeling research as an inverse problem. To illustrate the method, I collected calcium imaging data from the first two regions of the olfactory processing pathway of the fruit fly *Drosophila melanogaster*, the antennal lobe and the calix of the mushroom bodies. In each case, our method gives accurate predictions for large fractions of recorded glomeruli and neurons, and the inferred networks recover known features of the biological counterpart.

## INTRODUCTION

A model of the nervous system able to generate the behavior of the modeled organism remains a hard challenge. Brain function depends on a large number of variables, from topological features of the network to cellular and synaptic properties, and this makes it difficult to build a model of it from first principles. However, if the focus is on nervous system function (as opposed to the physical implementation of this function in biological brains), it should be possible to set this modeling research on a level that abstracts from a relevant part of this complexity. A prime example of this perspective is the application of deep learning to neuroscience, which has led to significant advances in modeling computations that range from sensory to motor functions^1–5^. The power of machine learning stems from optimization algorithms and from phrasing a particular question as an optimization problem, but I argue that there is a large amount of information that can be gathered from neuroscientific experiments that is not traditionally exploited in deep learning models. To this end, I propose to frame this modeling research as an inverse problem^6–10^, in order to take advantage of both optimization techniques and experimental neuroscience. In this context, the inverse problem is the problem of inferring an effective neural network given the recorded time series of neural activity. Data-driven research on dynamical systems has been extensively conducted in recent years, with various methods proposed to extract the dynamics of systems using their measured data^11,12^. However, reliable methods for the reconstruction of neural networks interactions from measurement data are a challenge due to the complexity of the system dynamics. *Drosophila melanogaster* olfactory processing pathway^13–27^ is ideal to test this approach as the relatively small number of neurons and the genetic toolbox available allow the recording of *in vivo* neural activity in both specific and comprehensive neural populations. The antennal lobe is the first stage of olfactory processing in this species. Olfactory receptor neurons (ORNs) in the antennae project their axons into the antennal lobe. There exist approximately 50 different types of receptor neurons and each class sends its axons to a single glomerulus, the functional sub-unit of the antennal lobe. In the glomeruli, ORNs synapse onto approximately 150 projection neurons (PNs) that send projections to the mushroom body and the lateral horn. Most PNs receive inputs from a single glomerulus^13^. There are also different types of local neurons (LNs) that have branches that are limited to the antennal lobes^16,17,21,22^. PNs synapse in an apparently random wiring process onto approximately 2000 Kenyon cells (KCs), the mushroom body intrinsic neurons^28^. Here I propose a combination of experimental recordings and a constrained regression algorithm to generate a partial effective model of the antennal lobe (AL) and the calix of the mushroom bodies (CX), as a proof of principle for a framework that should be possible to extend in a modular way to the whole brain.

## RESULTS

In order to infer an effective model of the first stages of the olfactory processing network of Drosophila, I presented a sequence of 21 olfactory stimuli (see Fig. 1b and Methods) to three groups of flies expressing GCamP6f in olfactory receptor neurons, projection neurons and Kenyon cells, targeted using Orco-, GH146- and OK107-GAL4 lines, respectively. The goal is to estimate the functions *f_i_*, *H_ij_* and *K_ij_* from the differential equations

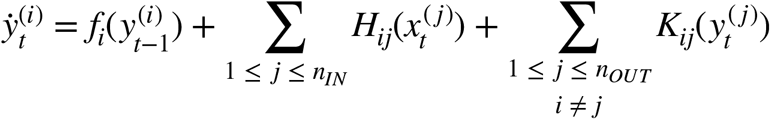

where 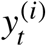 is the activity of the output unit *i* at time *t*, 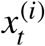 is the activity of the input unit *i* at time *t*, *f_i_* is the function that represents the dynamics of the isolated neuron, *H_ij_* are functions that describe the coupling between output unit *i* and input unit *j*, *K_ij_* are functions that describe the lateral coupling between output unit and output unit *j*, *n_IN_* is the number of input units and *n_OUT_* is the number of output units. In the first experiment (Fig. 2), ORNs have the role of input units and PNs of output units, while in the second experiment (Fig. 3) PNs are the input units to KCs. The inference algorithm to solve the inverse problem is a delicate procedure that is prone to overfitting and large prediction errors if left unconstrained, and particular care was required to obtain reasonably accurate predictions (see below and Methods for the details of the estimation process). Given the measured data, the network estimation can be formulated in general as a linear inverse problem for each output unit *i*^11,12^. At first the focus is on a simplified form of these equations by disregarding lateral interactions, *K_ij_* = 0, and by assuming linear functions, 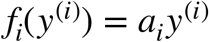 and 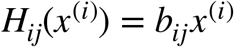 with *a_i_*, *b_ij_* ∈ **IR**. Solving the inverse problem means solving

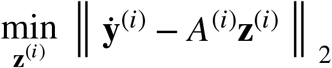

where 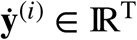 with *T* the total number of timepoints is the vector whose elements 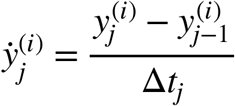 denote the finite difference approximation of the time derivative of output unit *i* at time *t_j_*, with Δ*t_j_* = *t_j_* − *t_j_*_−1_ being the data sampling time interval; 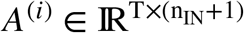 is the matrix defined by

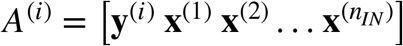

and 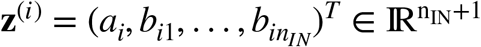 is the coefficients vector to be estimated for each output unit *i*. However, the straightforward implementation of this minimization problem leads to poor predictions, and the addition of supplementary inequality constraints on the parameters is required. Specifically, first an *l*_1_ penalty is imposed on the *b_ij_* coefficients to enforce a sparse solution; second, the decay time constants *τ_i_* = − 1/*a_i_* are required to be greater than 50 ms; third, the *b_ij_* coefficients were bounded to be excitatory connections, i.e. *b_ij_* ≥ 0 (see Methods).

**Fig. 1.**
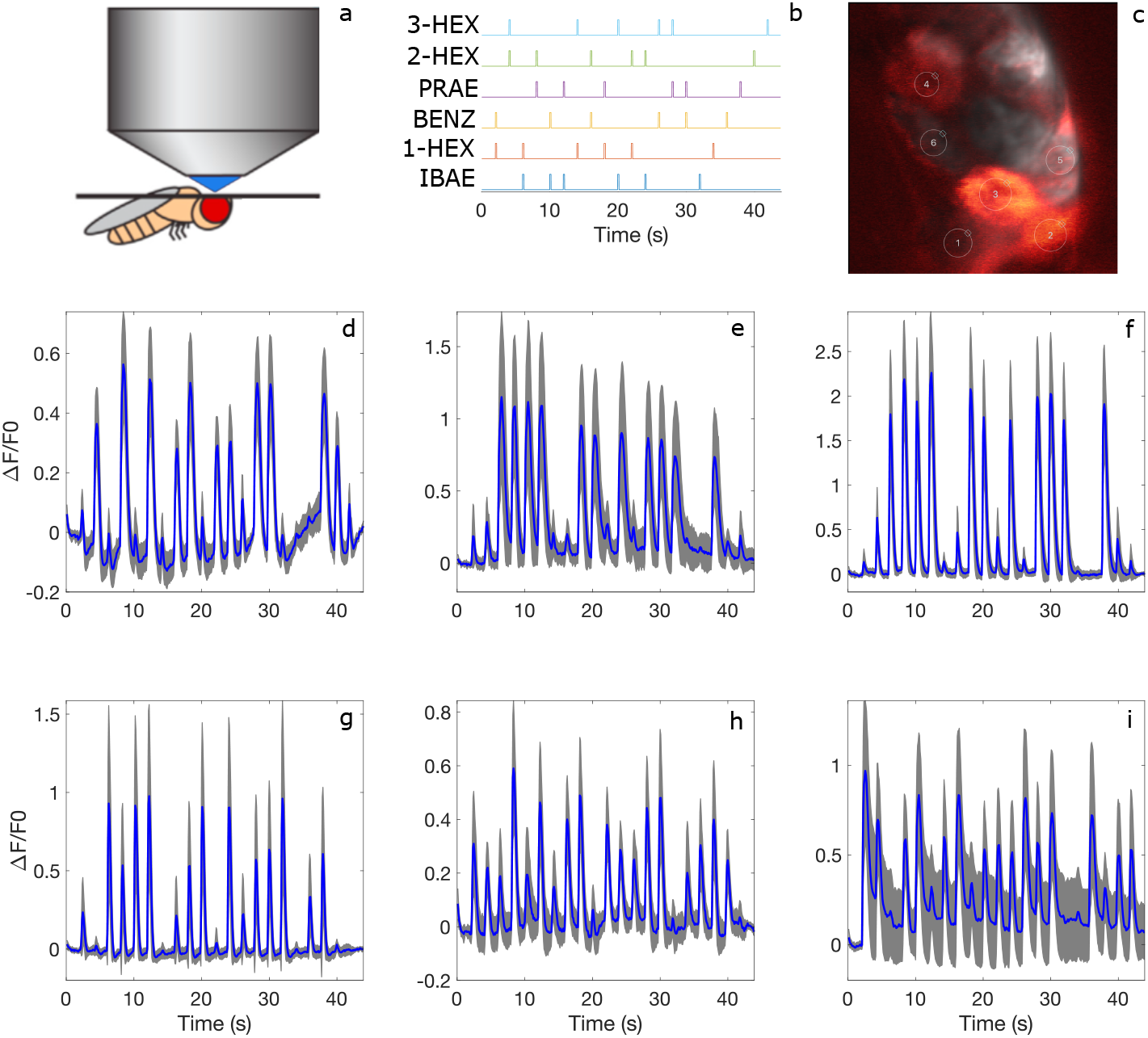
ORNs glomerular responses. **a**, Experimental setup. Restrained flies were presented with a sequence of olfactory stimuli while neural activity is monitored with calcium imaging. **b**, 200 ms stimuli were presented every 2 s, either as a single odor or as an odor pair. The voltage steps indicate the valve opening signals for the 6 odor channels. IBAE: isobutyl acetate; 1-HEX: 1-hexanol; BENZ: benzaldehyde; PRAE: propyl acetate; 2-HEX: 2-hexanol; 3-HEX: 3-hexanol. **c**, Time averaged fluorescence signals from a typical AL recording of ORNs axon terminals. ROIs of recorded glomeruli are overlayed. 1: DM1; 2: DM2; 3: DM3; 4: DC1; 5: DM5; 6: DL5. **d-i**, Trial averaged Δ*F*/*F*_0_ of the 6 recorded glomeruli, in the same order as in **c**. Standard deviation is displayed as grey shading. Data from *n* = 6 flies, 3 trials per fly. Each glomerulus was recorded in at least 3 flies.

**Fig. 2.**
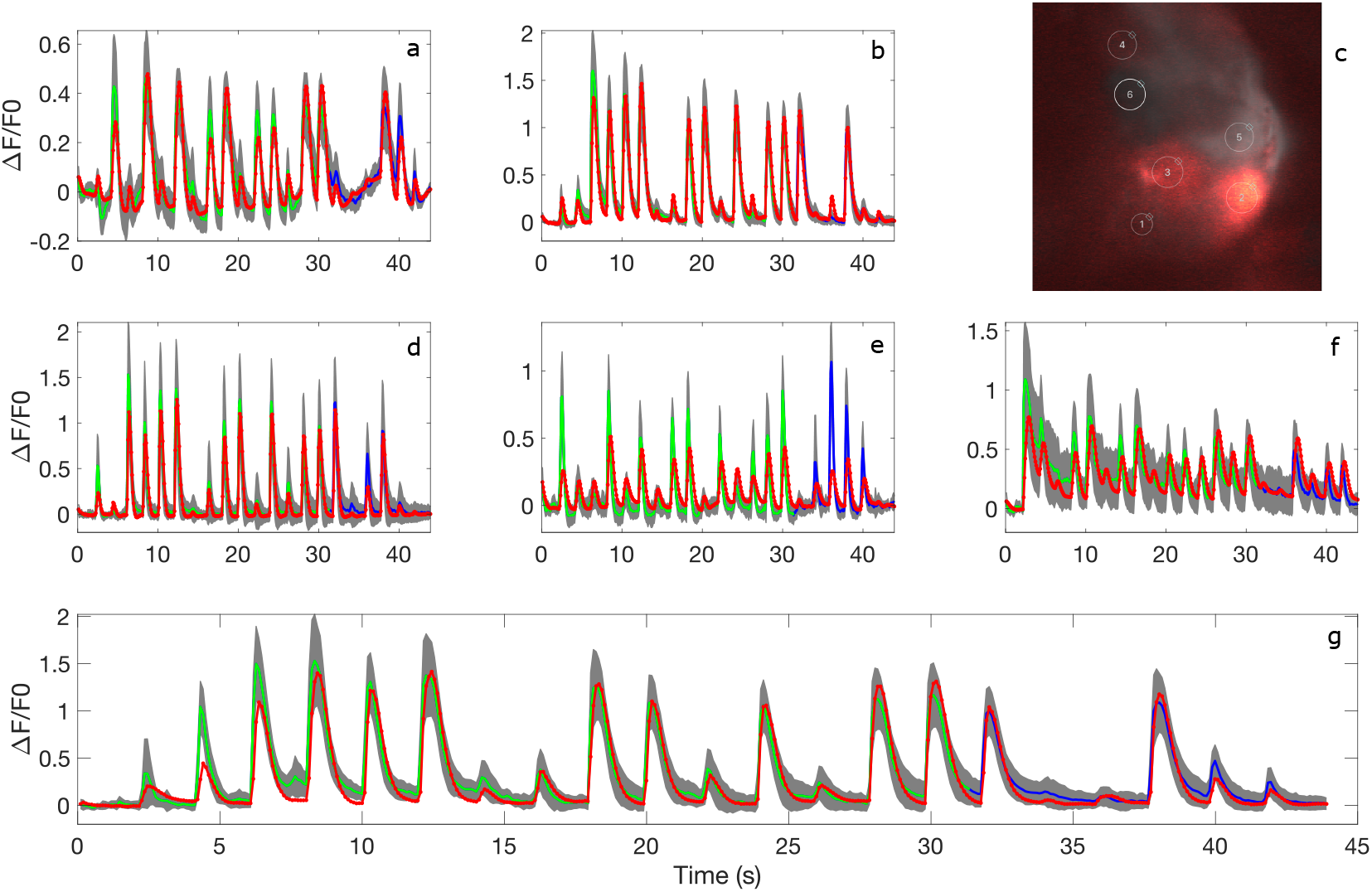
PNs glomerular responses and predictions of the inferred dynamical system. **a**, **b**, **d-g**, Trial averaged Δ*F*/*F*_0_ of the 6 recorded glomeruli. In green the segment used for training the model, in blue the data not presented to the inverse problem algorithm. Standard deviation is displayed as grey shading. In red, Δ*F*/*F*_0_ dynamics predicted by the inferred network. Data from *n* = 6 flies, 3 trials per fly. Each glomerulus was recorded in at least 3 flies. **a**, DM1, *E* = 0.67. **b**, DM2, *E* = 0.54. **d**, DC1, *E* = 0.51. **e**, DM5, *E* = 1.5. **f**, DL5, *E* = 0.76. **g**, DM3, *E* = 0.54. **c**, Time averaged fluorescence signals from a typical AL recording of PNs dendritic terminals. ROIs of recorded glomeruli are overlayed. 1: DM1; 2: DM2; 3: DM3; 4: DC1; 5: DM5; 6: DL5.

**Fig. 3.**
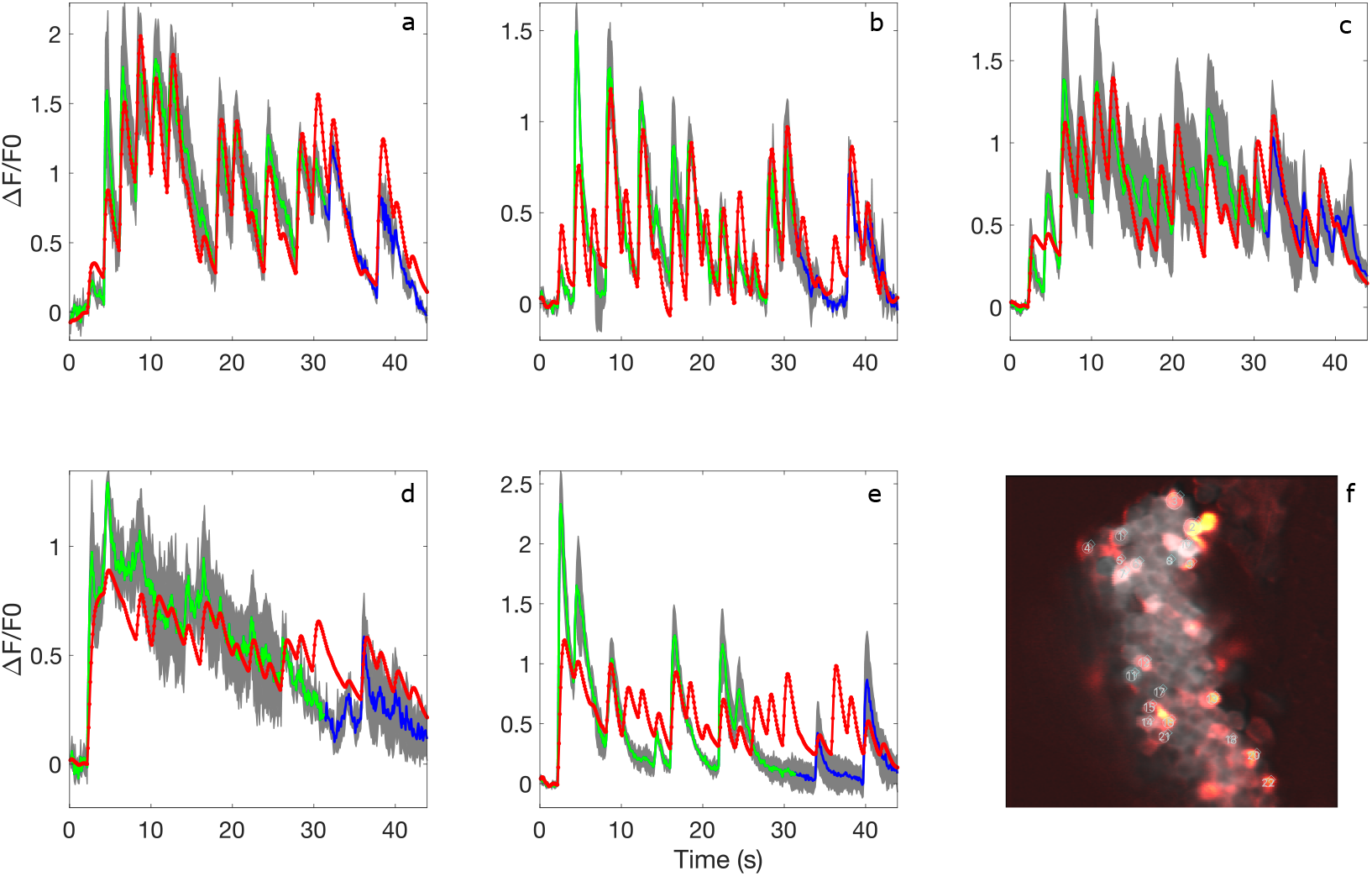
KCs somata responses and predictions of the inferred dynamical system. **a**-**e**, Trial averaged Δ*F*/*F*_0_ of 5 representative KCs. In green the segment used for training the model, in blue the data not presented to the inverse problem algorithm. Standard deviation is displayed as grey shading. In red, Δ*F*/*F*_0_ dynamics predicted by the inferred network. Data were pooled from *n* = 10 flies, 3 trials per fly. **a-c**, 3 cells with *Q* < 0.2. **d, e**, 2 cells with *Q* > 0.2. **i**, Time averaged fluorescence signals from a typical mushroom body recording of KCs somata. ROIs of recorded KCs are overlayed.

### Antennal lobe network

To infer the transformation from ORNs to PNs activities I recorded fluorescence changes in 6 glomeruli (Figs. 1c, 2c) identified based on their anatomical position and response properties. Given the anatomy of the antennal lobe, glomerular recordings should contain the same information as multiple recordings from ORNs expressing the same receptor, on one side, and of sister PNs (PNs synapsing onto the same glomerulus), on the other. I recorded Δ*F*/*F*_0_ traces from *n* = 6 Orco flies, 3 trials per fly, and averaged across trials and flies to obtain the responses in Fig. 1d-i. The same procedure was applied to GH146 flies (*n* = 6) to obtain the recorded traces in Fig. 2. These time series were then fed into our regression algorithm in order to infer an effective model for the ORN to PN transformation. The model was trained on the first 15 stimuli of the stimulation protocol and then tested on the whole sequence, with the red traces in Fig. 2 representing the predicted evolution of the output PN activity based on the experimentally recorded activity of the input ORNs. I argue that a true test to establish if the inferred network is sufficiently accurate will only be possible when a full sensorimotor transformation is modeled and the model is tested in the environment to assess whether it can generate the appropriate behavior. At this stage, however, I quantified the accuracy of the predictions by comparing the root mean square error (RMSE) between the prediction and the trial-averaged recorded trace to the mean RMSE between single trials and the trial-averaged recorded trace, computed on the test segment (blue segments in Fig. 2). A value of 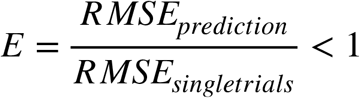 signifies a prediction that is accurate within the limits of inter-trial variability. For all glomeruli included in the analysis *E* < 1 except for DM5, for which *E* = 1.5 (Fig. 2e). The larger error in DM5 partly stems from the inaccurate prediction of the response to benzaldehyde (third peak of the test segment), but I was unable to identify the origin of this mismatch.

I also tested whether and how completely excluding from training one odor influenced the quality of the predictions. This corresponds to defining as training set all stimuli in the sequence that do not contain a given odor. The exclusion of one odor kept our results mostly unchanged, still maintaining *E* > 1 for DM5 and *E* < 1 for all other glomeruli, except in the following cases: for DM2, the exclusion of isobutyl acetate or propyl acetate raised the error to *E* = 1.44 and *E* = 1.54, respectively; for DM1, the exclusion of benzaldehyde or propyl acetate raised the error to *E* = 1.05 and *E* = 1.06, respectively; for DL5, the exclusion of benzaldehyde raised the error to *E* = 1.04 (data not shown). The larger prediction errors for DM2 when isobutyl acetate and propyl acetate are excluded from training is possibly due to the fact that this glomerulus responds strongly to these 2 odors, and their exclusion from training removes relevant information for the inference of accurate coupling parameters.

### Calix network

I then performed a similar experiment in order to model the transformation between PNs and KCs, without changing any of the parameters in the inverse problem algorithm. As input units I considered the same 6 glomerular responses recorded in GH146 flies shown in Fig. 2. I recorded *n* = 118 KCs pooled from *n* = 10 flies, 3 trials per fly. Predictions for 5 KCs that recapitulate the results for the whole population are displayed in Fig. 3.

Having only 3 trials per cell, prediction accuracy was quantified by evaluating *RMSE_pred_* in the test segment after normalizing each trace by the maximum of Δ*F*/*F*_0_ in that segment, *Q* = *RMSE_pred_* /*max*((Δ*F*/*F*_0_)_*test*_). This value was compared to the largest value of *Q* = 0.2 for the PNs predictions of Fig. 2, corresponding to the *Q* value for glomerulus DL5, and considered “accurate” any KC prediction with *Q* < 0.2. According to this metric, 37% of KCs were predicted well, and for these cells the predicted trajectory in neural state space largely recapitulates the trajectory for the trial averaged recorded activity (Video 1).

**Video 1.**
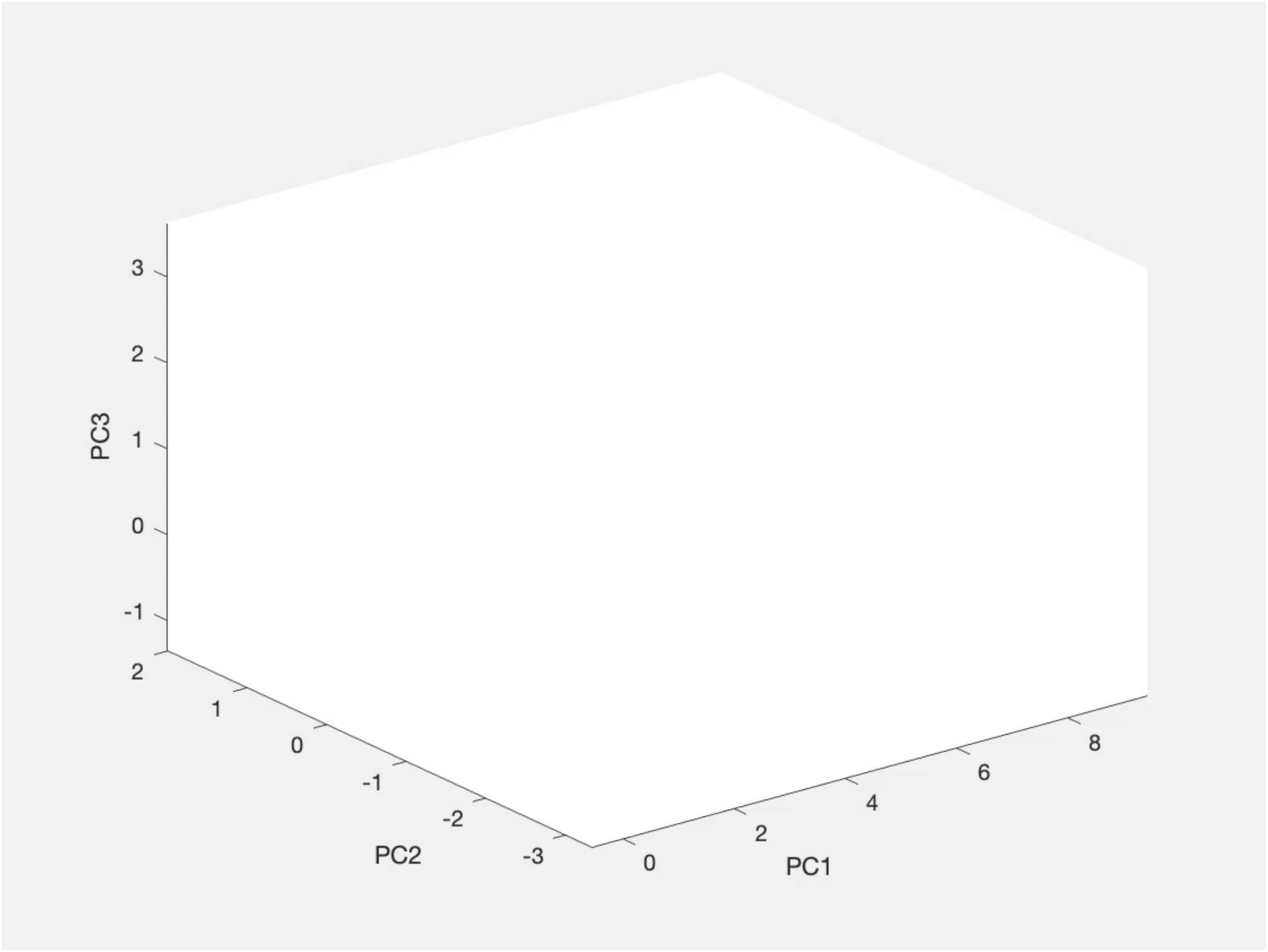
KC population trajectory. Neural trajectories in the test segment for KCs with *Q* < 0.2. Red: predicted activity; thick blue: trial averaged experimental activity; thin blue: single trial experimental activities.

Cells in Figs. 3a-c belong to this class, while Figs. 3d, 3e illustrate 2 examples of cells with *Q* > 0.2. Predictions were often poor in cells with slow decays and that responded strongly to just a few stimuli, as in Fig. 3e. It is also interesting to notice that the mismatch in the prediction of benzaldehyde alone (third peak of the test segment) was evident also in some KCs (Fig. 3b). Adding the possibility of lateral (excitatory and inhibitory) interactions between KCs by allowing *K_ij_* ≠ 0 did not appear to improve prediction quality (data not shown).

## Analysis of inferred effective networks

I argue that the primary goal of this modeling effort is to be able to predict the activity of the output units, irrespective of whether or not the model network has any similarity to the biological one^7^. At the same time, however, insights into the biological implementation of neural computations might prove very valuable. Our constrained regression algorithm was able to discover the known uniglomerular organization of PNs arborizations in the AL, as shown by the dominance of the diagonal terms in the inferred interaction matrix (Table 1). The decay time constants *τ_i_* = − 1/*a_i_* varied approximately between 100-400 ms.

**Table 1.** Inferred AL network. Each column represents the coupling between a given PN-side glomerulus and all ORN-side glomeruli plus the self-interaction coefficient *a* = − 1/*τ*. Red boxes highlight the interaction terms between PNs synapsing onto a given glomerulus and corresponding ORNs projecting to that same glomerulus.

| | $i = DM1$ | $i = DM2$ | $i = DM3$ | $i = DC1$ | $i = DM5$ | $i = DL5$ |
| --- | --- | --- | --- | --- | --- | --- |
| $a_i$ | -4.13 | -7.35 | -8.69 | -9.32 | -4.46 | -2.67 |
| $b_{i,DM1}$ | 3.32 | 0 | 0 | 0 | 0 | 0 |
| $b_{i,DM2}$ | 0 | 4.18 | 0.61 | 0 | 0 | 0 |
| $b_{i,DM3}$ | 0.14 | 0.95 | 5.02 | 1.71 | 0 | 0 |
| $b_{i,DC1}$ | 0 | 4.87 | 0 | 8.20 | 0.18 | 0.31 |
| $b_{i,DM5}$ | 0 | 0 | 0 | 0 | 4.40 | 0 |
| $b_{i,DL5}$ | 0.12 | 0 | 1.16 | 0.10 | 0 | 2.57 |

In the KCs experiment I limited the analysis to the cells whose activity was predicted well (*Q* < 0.2). Time constants for these cells were slower than for PNs glomerular predictions, ranging between 400 ms to 2.2 s. Moreover, the algorithm inferred a comparable number of connections onto KCs from each of the sister PNs groups, with similar average coupling strength onto KCs for all PN groups^28^ (Fig. 4). The exception is represented by sister PNs receiving input from DM2, which was always neglected as input to KCs. It is likely that this is an artifact of the regression algorithm: given the constraints imposed by *l*_1_ regularization and the fact that DM2 and DC1 responses to our stimulation protocol are very similar, the minimization process could assign the full contribution to the activity of a KC to only one of the 2 PN groups.

**Fig. 4.**
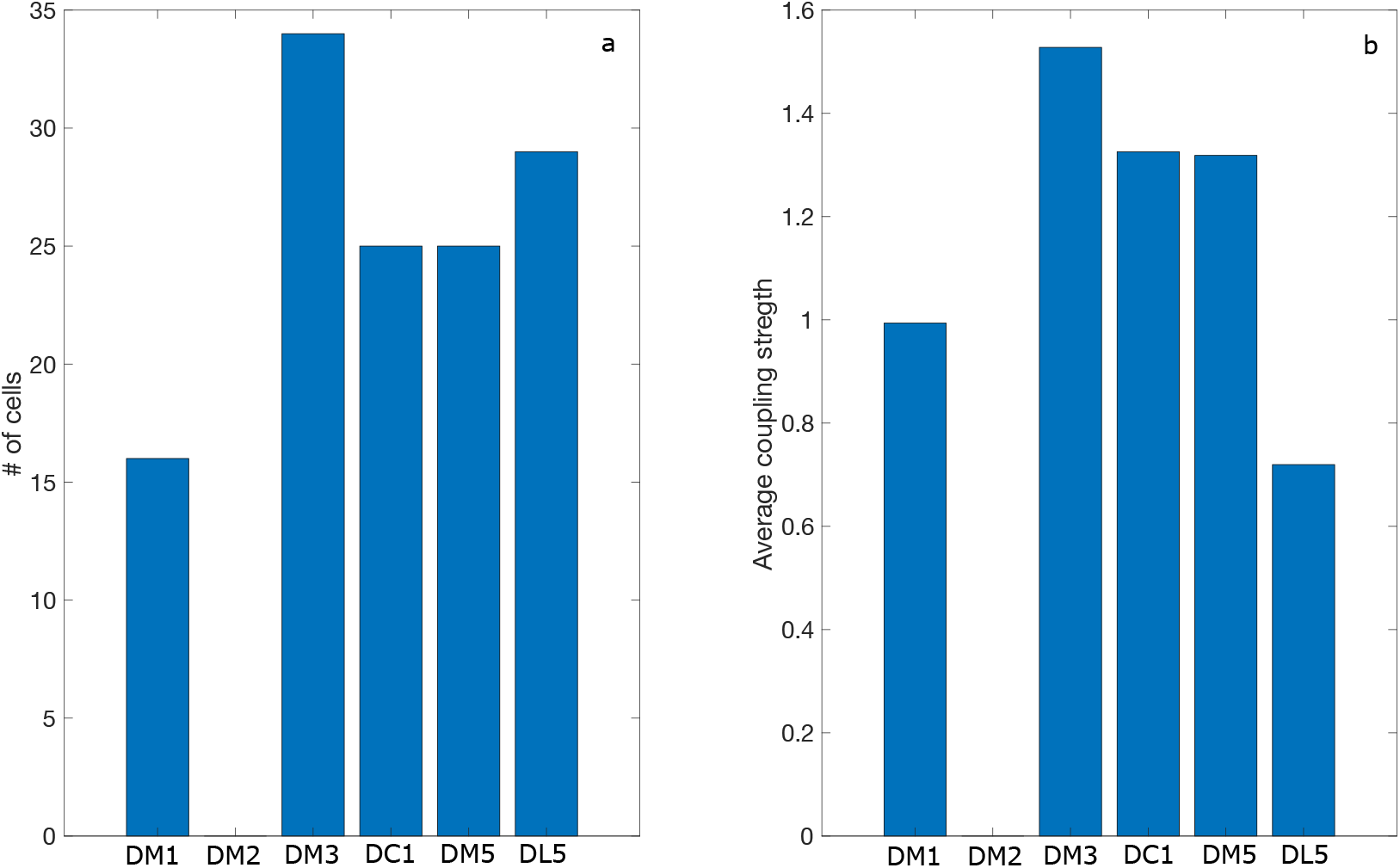
Properties of the inferred PN-KC interactions. **a**, Bar plot illustrating the number of Kenyon cells with positive coupling to the 6 PN groups. **b**, Average coupling strength to the whole KC population from the 6 PN groups.

## Discussion

Despite the success of connectomics projects^14,26^, the realization of working models of brain function is still an open issue. I adopt here a general approach for automated network inference in the framework of inverse problems. Ideally, the whole brain should be recorded simultaneously in the same animal. It is a difficult technical problem to achieve the required spatial and temporal resolutions even in *Drosophila*. However, I argue that, with the assumption that brain regions are functionally stereotypical across animals, it is possible to tackle the problem in a modular way, by recording from different areas in different animals and letting the regression algorithm generate a transformation between input and output units. This transformation might not be actually implemented in any animal, but would still represent a plausible functional network for that specific computation. The assumption, in other words, is that it is possible to use neural activity in a region of the brain of a specific animal as a complete basis to generate the responses of afferent regions in other organisms of the same species.

In the current version, the inferred dynamical system is static, i.e. the coupling terms do not change with time. A protocol was selected on purpose with short and separated stimuli that would minimize the effects of short-term synaptic plasticity mechanisms such as adaptation, in order to search for the “base”, or non-adapted, network. Once that is available, it is straightforward to add additional equations to the set differential equations modeling the system, in order to modulate the coupling functions between units (for example with known or hypothesized activity-related short-term plasticity). In this framework it is also not problematic to extend the model to incorporate any non-linear couplings, as described in ***Brunton SL, 2016***.

Predictions in the AL were reasonably accurate even without the addition of lateral interaction between PNs, which could also take into account the effects of the local interneurons. Either these lateral interactions were not strongly elicited by our stimulation protocol, or they mostly acted only at the ORNs level, for which the experimentally recorded activity was always used (so that the effects of the local neurons was already incorporated into the ORNs traces). In the MB, good predictions were obtained for 37% of recorded KCs. This could be due to the inadequacy of the regression algorithm itself, but it could still represents a notable result given that only olfactory inputs from 6 out of ~50 sister PNs groups were used and that KCs are known to receive input from different sources^26^. Next, it will also be important to analyze how this approach scales with the number of input units, as for example in the case of the transformation between KCs and mushroom body output neurons.

If successful, the perspective described can lead to the systematic development of a functional model of the fly brain. This would represent an important step towards the implementation of artificial systems with flexible and robust behavior.

## Materials and methods

### Fly strains

Flies were raised on cornmeal food at 25 ̊C and 60% humidity under controlled 12 hr light/dark cycles. All experiments were performed at room temperature (21 ̊C) on female flies that were 7 to 14 days old. Fly strains used were as follows: UAS-GCaMP6f, Orco-GAL4, GH146-GAL4, OK107-GAL4.

### Calcium imaging

The calcium fluorescent reporter GCaMP6f was used throughout the paper. Flies were briefly anesthetized on ice and fixed on a custom-built holder that left the antennae exposed to air. The proboscis was glued to the thorax with low-temperature-melting wax to reduce brain movement. The head capsule was opened and covered with Drosophila ringer (5 mM Hepes, 130 mM NaCl, 5 mM KCl, 2 mM MgCl2, 2 mM CaCl2, 36 mM sucrose, pH 7.3). For optical imaging, I used a LSM5 confocal microscope (Zeiss), with a Plan-Apochromat 20x/1 water-immersion objective (Zeiss). The excitation wavelength was 488 nm and images were recorded at 10 Hz.

### Odor delivery

The odor delivery system provides a constant total flow of approximately 1.2 liters per minute, with a primary and 6 secondary air streams of 1 and 0.03 liters per minute respectively. The 6 secondary streams correspond to the 6 odor channels and a solenoid valve switches them between air and odor modes. Delivered odors are continuously removed through a large suction tube. Odors were prepared at 10^−2^ concentration (volumetric ratio in mineral oil).

### Stimulation protocol

Odors were chosen to activate multiple glomeruli that are covered by both Orco and GH146 lines^27^ and to have weak surface interactions with the odor delivery system. The selected odors are isobutyl acetate, 1-hexanol, benzaldehyde, propyl acetate, 2-hexanol and 3-hexanol. The stimulation protocol was designed to minimize the effects of short-term plasticity mechanisms such as adaptation^19^, since I assumed time-independent interactions. Stimuli were presented for 200 ms each with a 2 s inter-stimulus interval, in a sequence of all possible pairs of odors (15 stimuli) plus the 6 single odors, 21 stimuli total for a duration of approximately 45 seconds. The full sequence is presented 3 times to each fly, with ~1 minute separation between trials.

### Image processing

All data were analyzed in MATLAB (Mathworks), except for calculation of the mean intensity of the regions of interest that was performed in custom python software. Images in Kenyon cells experiments were corrected for movement using NoRMCorre rigid correction^29^. Mean basal fluorescence (*F*_0_) was calculated by averaging the activity during the 2 s preceding first stimulus onset. Calcium responses were quantified as relative changes in fluorescence (*F*(*t*) − *F*_0_)/*F*_0_. A linear bleaching correction was applied to all trials showing decreased baseline fluorescence after the stimulation protocol by interpolating between the average fluorescence in the first second and average fluorescence in the last second.

### Population trajectories

PCA was applied to the 5*T* × *N* matrix containing the concatenated time series of the predicted, trial-averaged, and 3 single trial KC data, with T length of the time series and N total number of recorded KCs. The first 3 principal components are then retained and the 5 trajectories are projected onto this space.

### Constrained regression

The optimization problems 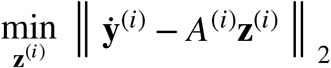 with inequality constraints on **z**^(*i*)^ was solved using MATLAB *lsqlin* function. The constraints can be leveraged to both add a regularization term and to introduce neuroscientific knowledge about the system into the optimization process. I argue that sparseness of the solution is an important feature, as only some of the recorded input units will usually be connected to a given output unit. Therefore, I chose to add an *l*_1_ penalty term ||**b**^(*i*)^ ||_1_ < *λ* to the minimization problem, where **b**^(*i*)^ is the vector with elements *b_ij_* defined by *H_ij_*(*x*^(*i*)^) = *b_ij_ x*^(*i*)^ (see main text). A way to make this constraint compatible with *lsqlin* is to decompose **b** into its negative and positive parts, **b** = **b**^+^ − **b**^−^, as the relation |**b**| = **b**^+^ + **b**^−^ handles the *l*_1_ constraint^30,31^. I also required the decay time constants 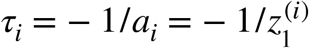 to be greater than 50 ms and bounded the *b_ij_* coefficients to be excitatory connections, i.e. **b**^(*i*)^ ≥ 0.

## ACKNOWLEDGEMENTS

I thank Giovanni Galizia, Carlotta Martelli and Alja Lüdke for discussions and comments on a previous version of the manuscript, Ajayrama Kumaraswamy for help with image processing software, and Alja Lüdke and Carlotta Martelli for technical assistance. This work was supported by funding from the University of Konstanz.

## Code availability

code is available at https://github.com/mstux/InverseProblem.

